# HOX13-dependent chromatin accessibility modulates the target repertoires of the HOX factors

**DOI:** 10.1101/789875

**Authors:** Ines Desanlis, Yacine Kherdjemil, Alexandre Mayran, Yasser Bouklouch, Claudia Gentile, Rushikesh Sheth, Rolf Zeller, Jacques Drouin, Marie Kmita

## Abstract

*Hox* genes encode essential transcription factors that control patterning during embryonic development. Distinct combinations of nested *Hox* expression domains establish cell and tissue identities^1–3^. Consequently, spatial or temporal de-regulation of *Hox* genes can cause severe alterations of the body plan^3^. While HOX factors have very similar DNA binding motifs, their binding specificity is, in part, mediated by co-factors^4–6^. Yet, the interplay between HOX binding specificities and the cellular context remains largely elusive. To gain insight into this question, we took advantage of developing limbs for which the differential expression of *Hox* genes is well-characterized^7^. We show that the transcription factors HOXA13 and HOXD13 (hereafter referred as HOX13) allow another HOX factor, HOXA11, to bind loci initially assumed to be HOX13-specific. Importantly, HOXA11 is unable to bind these loci in distal limbs lacking HOX13 function indicating that HOX13 modulates HOXA11 target repertoire. In addition, we find that the HOX13 factors implement the distal limb developmental program by triggering chromatin opening, a defining property of pioneer factors^8,9^. Finally, single cell analysis of chromatin accessibility reveals that HOX13 factors pioneer chromatin opening in a lineage specific manner. Together, our data uncover a new mechanism underlying HOX binding specificity, whereby tissue-specific variations in the target repertoire of HOX factors rely, at least in part, on HOX13-dependent chromatin accessibility.

---

The *Hox* family of developmental genes encoding transcription factors (TFs) have pivotal roles in organogenesis and patterning. Their differential expression along the main body axis establishes positional information that instructs cells about their fate, ultimately generating morphological diversity^1–3^. The wide-range of HOX-dependent genetic programs is intriguing considering that the various HOX TFs have similar DNA-binding properties^6^. Several studies uncovered the relevance of co-factors, notably the PBX/Exd family of TFs, in increasing DNA-binding specificity^4–6^. Analyses using micromass culture from primary mesenchymal limb bud cells revealed that group 11 and 13 *Hox* genes cluster in the same subgroup^10^ despite the fact that HOXA11/HOXD11 are required for the zeugopod (forearm) developmental program^11^ while HOXA13 and HOXD13 are mandatory for digit development^12^ (Fig. 1a). This raises the question of the extent to which HOX TF specificity may be modulated spatially/temporally during embryogenesis.

**Figure 1.**
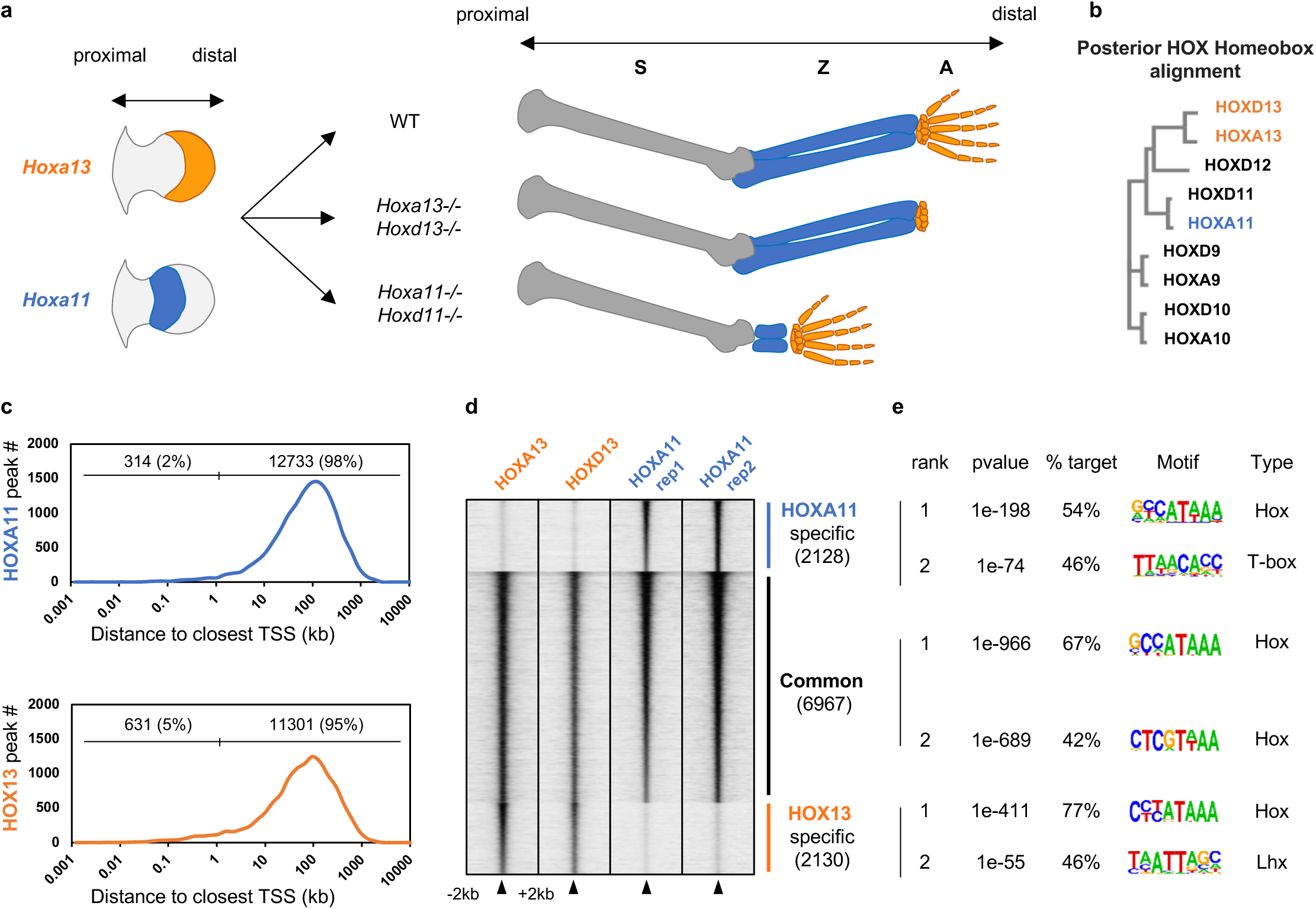
HOXA13 and HOXA11 bind common and specific genomic loci in developing mouse limb buds. (**a**) Schematics illustrating the mutually exclusive expression pattern of *Hoxa11* and *Hoxa13* in mouse limb buds (left) as well as the forelimb phenotype associated with the inactivation of *Hoxa/d11* and *Hoxa/d13* (right). (**b**) Alignment of 5’*Hoxa* and *Hoxd* homeobox sequences revealing the close relationship between *Hoxa/d11* and *Hoxa/d13* (**c**) Distance of the HOXA11 (blue) and HOXA13 (orange) bound loci to the closest TSS of all HOXA11 and HOXA13 peaks (p-value < 10^−5^) in E11.5 forelimb buds. (**d**) Heatmap showing ChIP-seq read density of HOXA11 (blue) and HOXA/D13 (orange) at all peaks for HOXA11 and HOXA13 with a p-value <10^−20^, in a 4kb window. Peaks are ranked based on p-value and according to the binding specificity. (**e**) Top scoring motif found by HOMER *de novo* motif analysis in a 200bp window around the peaks center of the three categories depicted in fig. 1d.

To address this question, we investigated the impact of the cellular context on the target repertoire of HOXA11 and HOX13, which are normally expressed in mutually exclusive domains in the developing limb but have highly similar homeodomain sequences (e.g. ref.10; Fig. 1b). We first analyzed the genome-wide binding of HOXA11 in wild type limb buds, at embryonic day 11.5 (E11.5), by performing Chromatin immunoprecipitation followed by high throughput sequencing (ChIP-seq) and compared the data with those previously reported for HOX13 (ref.13) (Fig. 1; Extended Data Fig.1). Reminiscent of the distribution of HOX13-bound loci in distal limbs^13^, the vast majority of HOXA11 ChIP-seq peaks are located at genomic regions distinct from transcriptional start site (TSS; Fig. 1c). While most peaks are common in both data sets, 23% of HOXA11 (i.e 2128) or HOXA13 (i.e 2130) targets are located at distinct loci (Fig. 1d) and *de novo* motif search uncovered slight differences in HOX motifs for common *versus* specific targets (Fig. 1e). We next tested whether the cellular environment could modulate the specificity distinguishing HOXA11 from HOXA13 binding sites. To this aim, we analyzed HOXA11 binding profile in *Prrx1:Cre;Rosa26*^*Hoxa11/Hoxa11*^ limb buds in which HOXA11 is ectopically expressed distally^14^ (*R26*^*A11d/A11d*^ in Fig. 2a). Compared to the wild type binding profile, we observed ectopic binding of HOXA11 in *R26*^*A11d/A11d*^ limb buds (Fig. 2b-c). Interestingly, distally expressed HOXA11 binds to loci, which, in wild type limbs, are specific to HOX13 and are enriched for the DNA motif associated with HOX13 specific sites (Fig. 2d, Extended Data Fig. 3a-c). To further assess whether these changes in HOXA11 binding are due to the presence of HOX13 or triggered by other factors specific to distal limb cells, we took advantage of the *Hox13-/-* mutant^12^ in which *Hoxa11* becomes distally expressed^13,15,16^. Analysis of HOXA11 genome-wide binding in *Hox13-/-* limbs revealed that all the ectopic HOXA11 binding sites observed in *R26*^*A11d/A11d*^ limbs were absent in *Hox13-/-* limbs (Fig. 2d), indicating that the ectopic binding of HOXA11 occurring in distal limbs requires HOX13 function. Together these results suggest that the binding specificity of HOXA11 and HOXA13 does not simply rely on distinct DNA motifs. Rather, our finding supports the idea that the primary driver for the distinct HOXA11 and HOXA13 binding profiles in wild type limb buds is related to their discrete expression domain, differentially expressed co-factors and possibly to chromatin accessibility.

**Figure 2:**
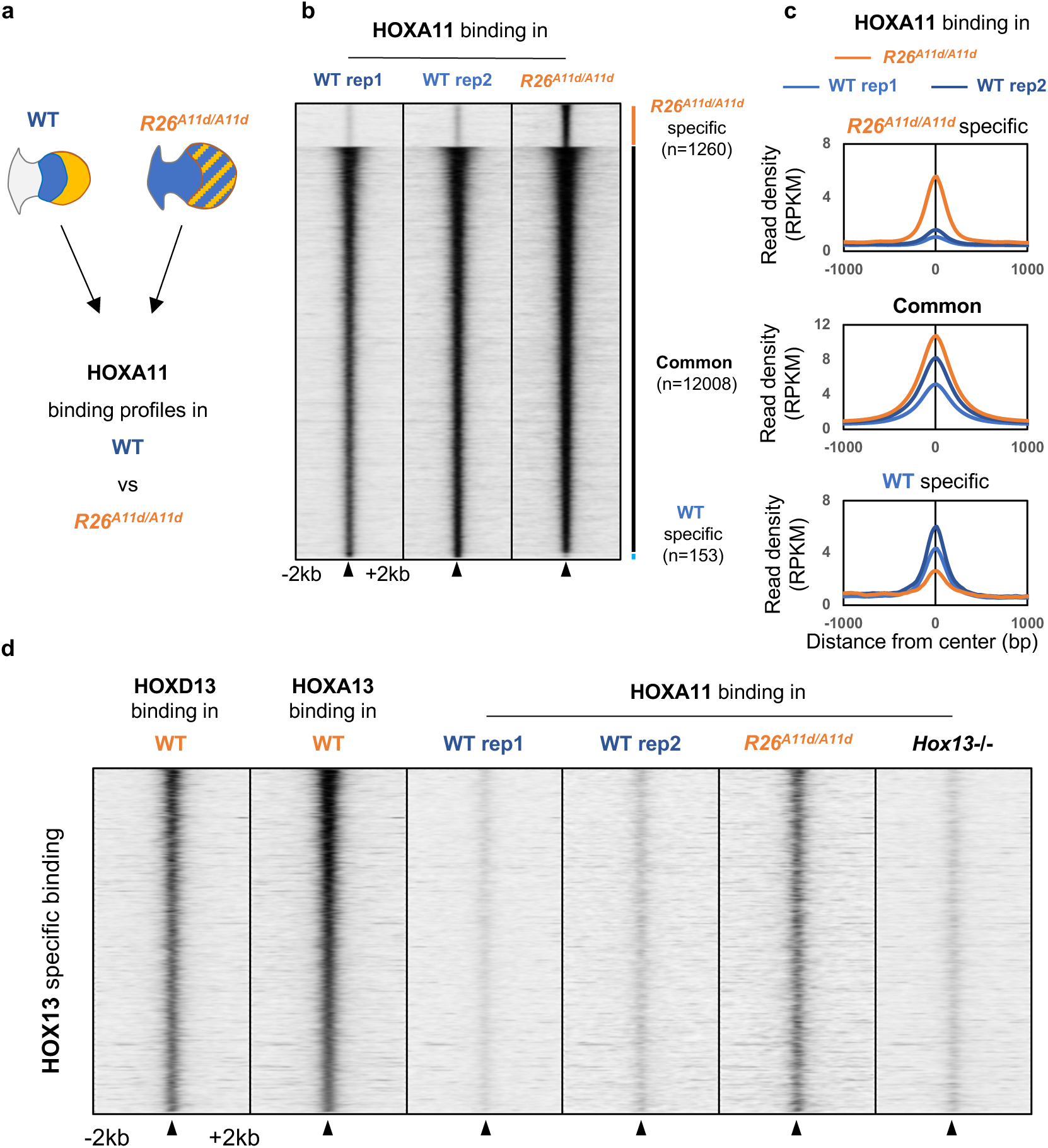
Loss of HOXA11-HOX13 specificity upon *Hoxa11* expression in distal limb buds. (**a**) Schematics showing the expression pattern of *Hoxa11* (blue) and *Hoxa13* (orange) in mouse limb bud from wild type and in *Prrx1Cre; Rosa*^*Hoxa11/Hoxa11*^ mouse (*R26*^*A11d/A11d*^). These were used to assess binding of HOXA11 associated with ectopic expression of *Hoxa11* in distal limbs. (**b**) Heatmaps showing a 4kb window of ChIP-seq read density for HOXA11 in wild type and *R26*^*A11d/A11d*^, at all HOXA11 peaks with a p-value <10^−20^. Peaks are ranked based on p-value and according to binding specificity. (**c**) Average profile showing ChIP-seq signal (RPKM) for HOXA11 in wild type (blue) and *R26*^*A11d/A11d*^ (brown) at the three categories depicted in fig. 3b. (**d**) Heatmaps showing a 4kb window of ChIP-seq read density for HOXA13/HOXD13 in wild type (left), HOXA11 in wild type (middle), *R26*^*A11d/A11d*^ and *Hox13-/-* E11.5 forelimb buds (right) at HOXA13 specific peaks with a p-value <10^−20^. Peaks are ranked based on p-value of HOXA13 binding.

**Figure 3.**
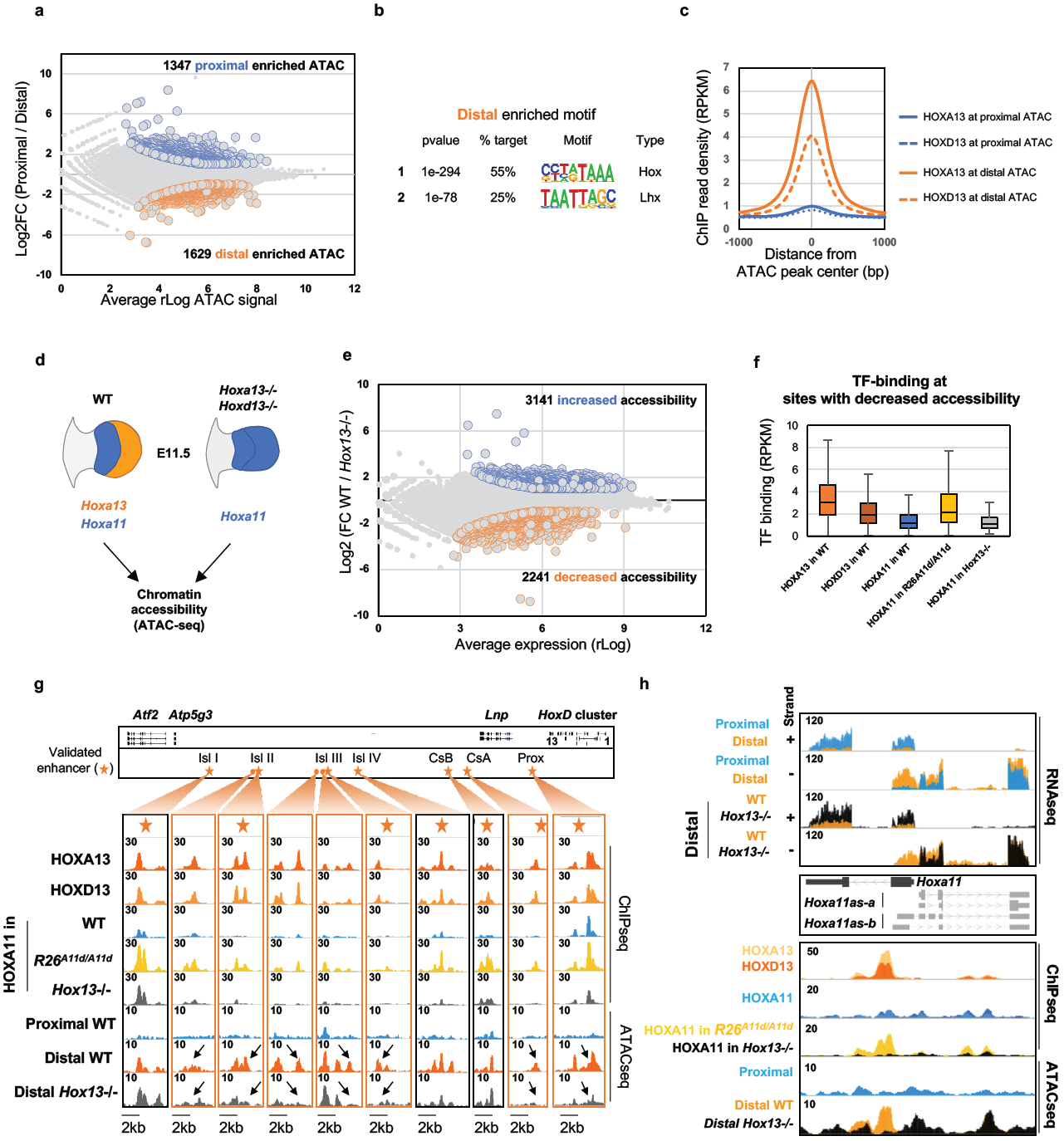
HOXA13 specific binding in wild type limbs correlates with HOX13-dependent chromatin accessibility. (**a**) MA Plot showing the average rLog (computed using deseq2) of ATAC signal (X axis) over the log2 fold change of proximal versus distal limb buds ATAC-seq peaks. Proximal-(blue) and distal-enriched accessibility (orange) are selected by the adjusted p-value <0.05, fold change > +/- 2). (**b**) Top scoring primary and secondary motifs found by HOMER *de novo* motif analysis in a 200bp window around the distally enriched peaks. (**c**) Average profile showing HOXA/D13 ChIP-seq signal (RPKM) at proximal (blue) and distal (orange) enriched ATAC peaks. (**d**) Scheme showing the expression pattern of *Hoxa11* (blue) and *Hoxa13* (orange) in wild type and *Hox13-/-* E11.5 forelimb buds. (**e**) MA Plot showing the average rLog (computed using deseq2) of ATAC signal (X axis) over the log2 fold change of wild type versus *Hox13*-/-distal E11.5 limb buds ATAC-seq peaks. Wild type (blue) and *Hox13*-/-(orange) enriched accessibility are selected by the adjusted p-value <0.05, fold change > +/- 2). (**f**) Boxplot showing the ChIP-seq signal read density (RPKM) for HOXA/D13, and HOXA11 in wild type, in *R26*^*A11d/A11d*^ and in *Hox13-/-* E11.5 limb buds over a 200bp window at sites with decreased accessibility. Center lines show medians; box limits indicate the twenty fifth and seventy-fifth percentiles; whiskers extend to 1.5 times the interquartile range from the twenty-fifth to seventy-fifth percentiles. pvalue were calculated using unpaired two-tailed student t-test. (**g**) Scheme of the genomic region encompassing previously validated enhancers (orange stars) responsible for *HoxD* distal limb expression (upper panel). Genome browser view (IGV, bottom) showing the intensity (reads per millions) of HOXA13, HOXD13 and HOXA11 ChIP-seq as well as ATAC-seq signals in wild type proximal and distal limb bud as well as in *Hox13*-/-distal limb bud. Loci with reduced chromatin accessibility upon *Hox13* inactivation are orange-framed and the arrow points at the site with loss/decreased accessibility. Note the loci with HOX13 ChIP-seq peaks, showing distal-specific, HOX13-dependent chromatin accessibility (orange ovals), which possibly coincide to distal limb-specific enhancers. (**h**) Genome browser view (IGV) of overlaid strand specific RNA-seq at the *Hoxa11* locus in the indicated tissues showing the mutually exclusive expression of *Hoxa11* and the antisense transcripts overlapping with *Hoxa11* exon1 (upper panel). Genome browser view (IGV) of the ChIP-seq and ATAC-seq data at the *Hoxa11* locus in wild type, *R26*^*A11d/A11d*^ and *Hoxa13-/-* limb buds. Note the HOX13-dependent chromatin accessibility at the enhancer driving *Hoxa11* antisense transcription and the ectopic HOXA11 binding (yellow) when *Hoxa11* is expressed distally, which is lost (black) upon *Hox13* inactivation.

To assess whether chromatin accessibility may influence HOXA11 and HOXA13 binding, we performed ATAC-seq (Assay for Transposase Accessible Chromatin followed by high-throughput sequencing^17^) on proximal and distal wild type limb buds. We identified 1629 sites showing stronger accessibility in distal limb buds (Fig. 3a, Extended Data Fig. 4a). These sites tend to be associated with genes involved in digit morphogenesis (Extended Data Fig. 4c-g) and are enriched for the previously identified HOX13 motif (Fig. 3b). Accordingly, HOX13 binds these distally accessible sites but not the proximal-specific accessible sites (Fig. 3c). Previous analysis of HOX13-dependent gene regulation revealed that the transition from the early to the distal/late limb developmental program relies on HOX13 function and it was proposed that such switch in developmental program could be associated with a potential pioneer activity of the HOX13 TFs^13^. The finding that cellular environment modulates HOXA11 binding in a HOX13-dependent manner (Fig. 2) thus prompted us to test whether the ectopic HOXA11 binding at loci normally specifically bound by HOX13 could originate from changes in chromatin accessibility mediated by HOX13 TFs. We thus performed ATAC-seq on wild type and *Hox13-/-* distal limbs (Fig. 3d, Extended Data Fig. 5a). We identified substantial changes in chromatin accessibility (Fig. 3e), and interestingly, loci with decreased accessibility were enriched for HOX13 binding in the wild type distal limb (Extended Data Fig. 5b). In contrast, loci with increased accessibility in *Hox13-/-* distal limbs did not overlap with HOX13 ChIP-seq peaks identified in the wild type context, suggesting that this gain in chromatin accessibility is an indirect outcome of *Hox13* inactivation (Extended Data Fig. 5b). We then analyzed the relationship between HOXA11 and HOX13, at sites loosing accessibility in *Hox13-/-* distal limbs. While HOXA11 and HOXA13 binding are correlated (R^2^=0.57) at sites with unchanged accessibility, HOX13-dependent accessibility shows an unambiguous preference for HOXA13-bound loci and a low correlation with HOXA11 (R^2^=0.18) (Extended Data Fig. 5e). Importantly, we found that expression of HOXA11 in distal limb cells leads to its ectopic binding at loci whose chromatin accessibility relies on HOX13 (Fig. 3f). Together these results suggest that HOX13-dependent chromatin accessibility allows HOXA11 binding at loci that are HOX13-specific in wild type limb, i.e. in a context where *Hox13* and *Hoxa11* are expressed in mutually exclusive domains.

Among the genomic regions where HOX13-dependent chromatin accessibility allows for HOXA11 binding, we found several previously characterized enhancers. For instance, several enhancers controlling *HoxD* gene expression in distal limb^18–20^ were identified as loci with ectopic HOXA11 binding (Fig. 3g). The HOX13-dependent chromatin accessibility is also consistent with the loss of transcriptional activity previously reported at these enhancers in *Hox13-/-* limbs^13,21^. Similarly, HOXA11 binding dependent on HOX13 is observed at the enhancer driving *Hoxa11* antisense transcription (Fig. 3h), whose function ultimately prevents distal expression of *Hoxa11*^14^. We next tested whether HOX13-dependent chromatin accessibility could be modulated by cell identity. To this aim, we performed single-cell ATAC-seq using wild type and *Hox13-/-* E11.5 forelimb buds (Extended Fig. 6). Cell clustering was based on the similarity of their genome-wide chromatin accessibility and the resulting clusters (Fig. 4a) were annotated using aggregated accessibility profiles of several marker genes (Extended Fig. 7). Consistent with *Hox13* being expressed in the limb mesenchyme, we found that clusters present in wild type and absent in *Hox13-/-* limbs (and vice-versa) are restricted to mesenchymal cells (Fig. 4b-c). More specifically, cluster 2 is completely lost upon *Hox13* inactivation and cluster 6 and 7 are drastically reduced (Fig. 4b), confirming the loss of chromatin accessibility uncovered from the bulk ATAC-seq analysis. In addition, occurrence of a *Hox13-/-* specific cluster (cluster 3, Fig. 4c) suggested that chromatin accessibility did not switch to a proximal limb profile but may correspond to an interrupted transition of progenitor cells towards specified distal limb cells. Analysis of the aggregated cluster ATAC-seq signal confirmed cell type specific loss of chromatin accessibility at HOX13 target sites. While our scATAC-seq data was consistent with the results obtained from bulk ATAC-seq assay, it also revealed additional loci dependent on HOX13 for their accessibility. For instance, the enhancer driving *Bmp2* expression in distal limb^22^ only showed reduced accessibility in bulk ATAC-seq (Extended Data Fig. 5g) while the scATAC-seq revealed an unambiguous loss specifically in clusters corresponding to distal mesenchyme (Fig. 4d, first column). Our data also suggested that the region previously identified as the *bmp2* distal limb enhancer likely encompasses two adjacent enhancers, with only one showing HOX13 dependency for its accessibility. Consistent with our results, *Bmp2* expression in *Hox13-/-* limbs is lost distally while unaffected in the proximal limb where *Hox13* is normally not expressed (Fig. 4e). Similarly, loss of chromatin accessibility in *Hox13-/-* limbs were observed at putative enhancers located in the neighborhood of genes expressed in distal limb buds (Fig. 4d) as well as at the previously identified enhancer driving *Hoxa11* antisense transcription distally (Fig. 4d, right column). Of note, ‘non-mesenchyme’ clusters were unaffected by *Hox13* inactivation (Fig. 4a), consistent with *Hox13* not being expressed in these cells.

**Figure 4:**
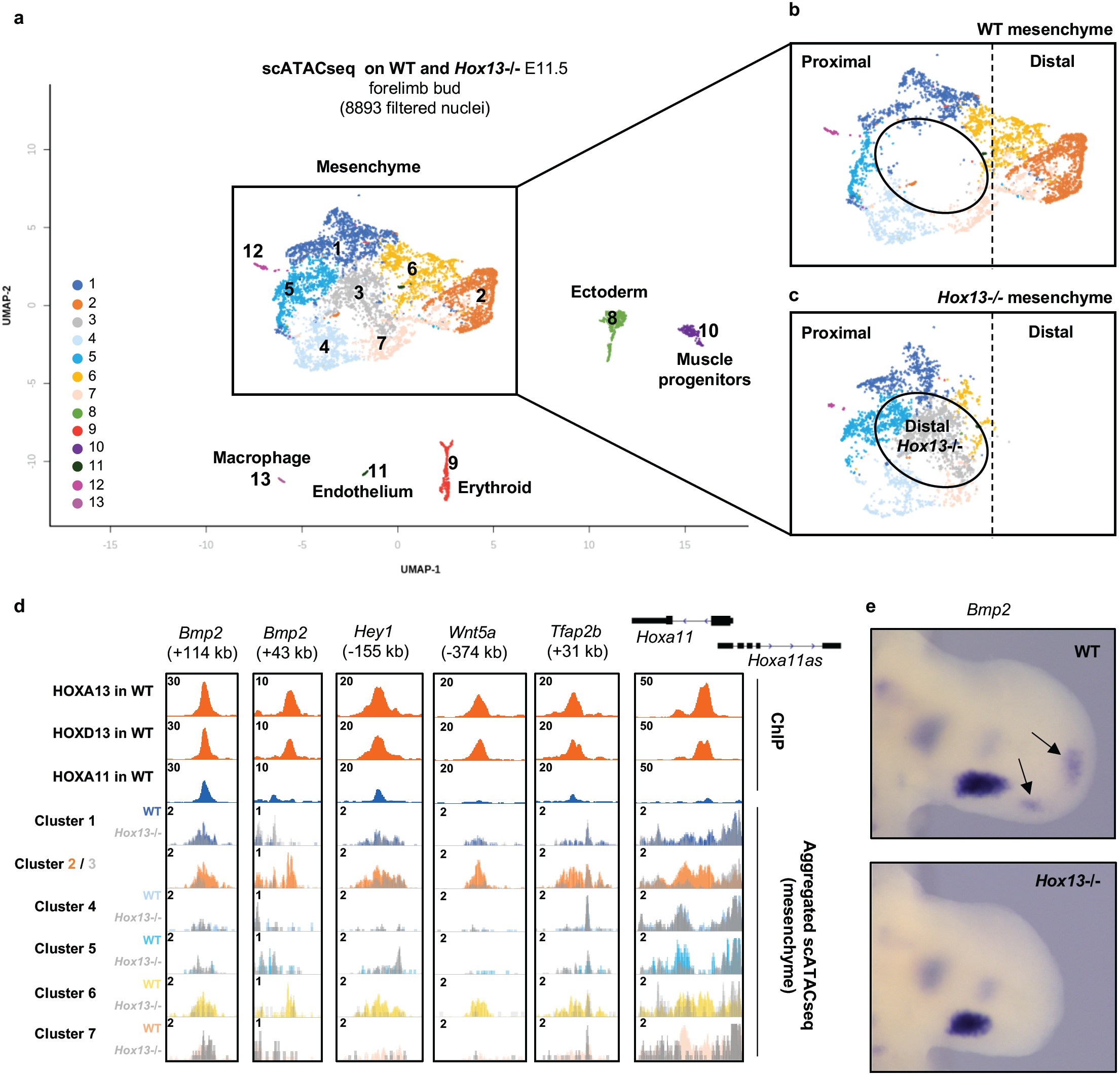
Single-cell ATAC-seq analysis reveals specific clusters with modified chromatin accessibility upon *Hox13* inactivation. (**a**) Dimensionality reduction map (UMAP) representation of the distribution of the various clusters identified using single cell ATAC-seq. Chromatin accessibility at promoter of genes known to be exclusively expressed in a specific cell population was used to assign cell identity to the clusters. (**b, c**) Magnification of mesenchymal clusters in wild type (top) and *Hox13-/-* (bottom). Cluster 2 (orange) is entirely lost in the *Hox13-/-* limb bud and cluster 6 (yellow) and 7 (light pink) are drastically reduced, while cluster 3 (grey) is specific to *Hox13-/-* limb buds. Note that clusters corresponding to non-mesenchymal cells (8 to 13) are unaffected by *Hox13* inactivation. (**d**) Genome browser view (IGV) of a subset of previously identified distal limb enhancers or potential distal limb enhancers associated with genes expressed in distal limb buds. ATAC-seq peaks in cluster 2 (orange) and 3 (grey) are superimposed to highlight the loss of accessibility upon *Hox13* inactivation. (**e**) Whole-mount *in situ* hybridization of *Bmp2* in wild type and *Hox13*-/-E11.5 limb buds. The arrows point to the two spots of distal *Bmp2* expression lost in *Hox13*-/-.

Together, these data provide evidence that HOX13 has the capacity of triggering the switch from a ‘closed’ chromatin conformation to an accessible one and allows the binding of other TFs. Whether this functional characteristic is specific to HOX13 or a common feature of the HOX family of TFs remains to be investigated but we propose that pioneer activity of HOX TFs is likely key to HOX-dependent cell fate and patterning events. Importantly, the evidence that HOX13 modifies the HOXA11 genome-wide binding repertoire reveals a mechanism that contributes to defining the tissue-specific spectrum of HOX targets. This mechanism does not belittle the importance of HOX cofactors as *de novo* motif analysis at loci with HOX13-dependent chromatin accessibility did not reveal an obvious motif distinguishing these loci from other HOX13 targets (Fig. 1e). Yet, the fact that a subset of HOX13 targets remain in an accessible chromatin state in *Hox13-/-* limb does not exclude the possibility that these targets could be switched to an accessible state by HOX13 in other cellular/tissue environments. Interestingly, a recent study investigating the *in vivo* HOX binding specificity at the *vvl1+2* enhancer in *Drosophila melanogaster* suggests that HOX binding specificity relies on a combination of three parameters: binding specificity influenced by HOX cofactor(s), HOX monomers and interaction with collaborator proteins^23^. Our work reveals that the HOX13 pioneer activity, and possibly that of the other HOX factors, is one of the key parameters that specifies the HOX target repertoires.

The impact of the HOX13 TFs on chromatin accessibility at *HoxD* enhancers raises the possibility that it contributes to the dynamic expansion of the *HoxD* expression domain during the progression of distal limb bud development. Interestingly, it was previously proposed that the increase/expansion of the *Hoxd13* expression domain might have been a driver for the origin of the novel endoskeletal elements characterizing the fin-to-limb transition^24^. In this view, the HOX13-dependent chromatin accessibility identified here for a subset of the *HoxD* enhancers might have contributed to the molecular evolution that allowed the emergence of digits. In addition, the HOX13-dependent chromatin accessibility at the enhancer driving antisense transcription at the *Hoxa11* locus, which is required to prevent distal *Hoxa11* expression and polydactyly^14,25^, supports a possible role of HOX13 pioneer activity in the molecular network that underlies the evolution of the pentadactyl state characterizing most modern tetrapods. Consistent with this view, we found that the enhancer driving *Hoxa11* antisense transcription is in an accessible state in the presumptive digit domain of chicken limb buds while it is in a ‘closed’ configuration in early limb buds, when *Hox13* genes are not expressed (Extended data Fig.9). This result corroborates the findings in wild type and mutant mouse limb buds and points to a conserved role of the HOX13 TFs in regulating chromatin accessibility during both mouse and chicken embryonic development. Based on these data, we propose that HOX13 pioneer activity could have been a key feature contributing to the emergence of digits and subsequent refinement of digit number in modern tetrapods.

## ACKNOWLEDGEMENTS

We are particularly grateful to lab members for insightful discussions and sharing reagents. This work was supported by the Canadian Institute for Health Research (MOP-115127 and -126110 to M.K and FRN-154297 to J.D.) and ERC advanced grant INTEGRAL (ID 695032) to R.Z. Bioinformatics analyses were enabled in part by support provided by Calcul Quebec (www.calculquebec.ca) and Compute Canada (www.computecanada.ca). Y.K. was supported by a fellowship from the Molecular Biology program of the Université de Montréal and the IRCM fellowship Michel-Bélanger. A.M. was supported by an IRCM challenge fellowship, C.G. was supported by the Jacques Gauthier IRCM fellowship, I.D. is supported by the IRCM-Jean Coutu fellowship.

## AUTHOR CONTRIBUTIONS

Y.K., A.M. and M.K. conceived the study. I.D., Y.K., A.M., designed the experiments. I.D. and Y.K., with the help of C.G. and A.M. conducted the experiments. R.S. and R.Z. provided the data shown in Extended. Fig. 9. A.M., with the help of I.D. conducted the bio-informatic analyses of all bulk datasets. Y.B. performed bioinformatics analyses of the single cell ATAC-seq data set. I.D., Y.K., A.M., and M.K. analyzed and interpreted the data. M.K. wrote the paper together with A.M., I.D., and Y.K. All authors commented on the manuscript.

## AUTHOR INFORMATION

The authors declare no competing financial interests.

Correspondence and requests for materials should be addressed to

## METHODS (ONLINE-ONLY)

### Mouse lines

*Hoxa13null (Hoxa13*^*Str*^*), Hoxd13null (Hoxd13*^*lacZ*^*), Rosa*^*Hoxa11*^ mouse lines were previously described^12,14,26^. All mice were maintained in mixed background (C57BL/6 × 129). Noon of the day of the vaginal plug was considered as E0.5. Mice and embryos were genotyped by PCR using genomic DNA extracted from tail biopsy specimens and yolk sacs, respectively. Mice work at the Institut de Recherches Cliniques de Montréal (IRCM) was reviewed and approved by the IRCM animal care committee (protocols 2015-14 and 2017-10).

### Chromatin Immunoprecipitation and Sequencing

HOXA11 ChIP was performed in forelimb buds of CD1 (wild type) and *Prx1Cre; Rosa*^*Hoxa11/Hoxa11*^ *(R26*^*A11d/A11d*^) mice at E11.5 in the same conditions as previously described for HOX13 ChIP^13^. Chromatin was cross-linked using a combination of disuccinimidyl glutarate (DSG) and formaldehyde and sonicated using Fisher Scientific, Model 100 sonic dismembrator to obtain fragments between 100-600 bp. Protein A and Protein G Dynabeads (Invitrogen) were incubated for 6 hours at 4°C with 5ug HOXA11 (SAB1304728, Sigma) antibody. The chromatin was coupled to the beads overnight at 4°C. The immunoprecipitated samples were then sequentially washed in low salt (1% Triton, 0,1% SDS, 150 mM NaCl, 20 mM Tris (pH8), 2 mM EDTA), high salt (1% Triton, 0,1% SDS, 500 mM NaCl, 20 mM Tris pH8, 2 mM EDTA), LiCl (1% NP-40, 250 mM LiCl, 10 mM Tris (pH8), 1 mM EDTA) and TE buffer (50 mM NaCl, 10 mM Tris (pH8), 1 mM EDTA). The DNA was then purified on QIAquick columns (Qiagen). Library and flow cells were prepared by the IRCM Molecular Biology Core Facility according to Illumina’s recommendations and sequenced on Illumina Hiseq 2500 in a 50 cycles paired-end configuration.

### ATAC-seq

Dissection of proximal and distal of forelimb buds from wild type embryos and distal forelimb buds of *Hox13*^*-/-*^ embryos^13^ were performed at E11.5. All samples for ATAC-seq were processed as previously described^27^. Briefly, 50 000 cells were washed in PBS and incubated on ice for 30 minute in a hypotonic cell lysis buffer (0.1% ^w^/_v_ sodium citrate tribasic dehydrate and 0.1% ^v^/_v_ Triton X100) centrifugated (5 minutes at 2000g at 4°C). Cells were then incubated 30 minutes on ice in cell lysis buffer (10mM Tris-HCl, pH7.4, 10mM NaCl, 3mM MgCl_2_, 0.1% ^v^/_v_ IGEPAL CA-630. After centrifugation (5 minutes at 2000g at 4°C) the resulting pellet of nuclei was re-suspended in transposase Master Mix (1.25 µl 10x TD buffer, 5 µl H_2_O and 6.5 µl of Tn5: Illumina Nextera Kit; FC-121-1031) and incubated for 30 minutes at 37°C. Samples were purified using MinElute PCR purification column (Qiagen). The eluted DNA was enriched and barcoded for multiplexing of samples using Nextera barcodes by PCR using Phusion kit. The library was recovered with GeneRead Purification columns. Samples were then evaluated by qPCR to test for the enrichment of open regions and sequenced on Illumina Hiseq 2500 with 50bp or paired end reads, according to Illumina’s recommendation.

### Single cell ATAC-seq

Dissection of forelimb buds from wild type embryos and *Hox13-/-* embryos^13^ were performed at E11.5. Forelimbs were dissected in cold 1XPBS, after centrifugation at 300 rcf for 5 minutes at 4°C the forelimbs were incubated in dissociation buffer (450 µl of 0.25% Trypsine/EDTA (GIBCO), 50 µl 10% BSA, 1 µl DNAseI (NEB)) for 10 minutes at 37°C. After incubation limb cells were gently mixed by pipetting up and down 10-15 times until they were dissociated, then 10% final FBS was added. Dissociated cells were filtered using a cell strainer (40 µm Nylon, BD Falcon) and after centrifugation at 300 rcf for 5 minutes at 4°C the resulting pellet of cells was resuspended in 1XPBS, 0.1%BSA. Dissociated cells were counted and assessed for cell viability using 0.4% Trypan blue. After centrifugation at 300 rcf for 5 minutes at 4°C, the cell pellet was resuspended in 100 µl of ice-cold lysis buffer (10mM Tris-HCl, pH7.4, 10mM NaCl, 3mM MgCl2, 0.1% Tween-20, 0.1% NP-40, 0.01% Digitonin, 1% BSA) and incubated for 5 minutes on ice. Then 1 ml of wash buffer (10mM Tris-HCl, pH7.4, 10 mM NaCl, 3mM MgCl2, 0.1% Tween-20, 1% BSA) was added to the cells. After centrifugation at 300 rcf for 5 minutes at 4°C, the pellet of nuclei was re-suspended in diluted nuclei buffer in order to have 3500 nuclei/µl and processed using Chromium Single Cell ATAC Reagent Kits (10x Genomics, Pleasanton, CA) following the manufacturer recommendation. Briefly, nuclei were incubated in a Transposition Mix that includes a Transposase that preferentially fragments the DNA in open regions of the chromatin and simultaneously, adapter sequences were added to the ends of the DNA fragments. GEMs were then generated, using partitioning oil, in order to contain a single nucleus and a Master Mix to produce 10x barcoded single-stranded DNA after thermal cycling. The Chromium Single Cell ATAC Library was amplified and double sided size selected. Samples were controlled at multiple steps during the procedure by running on BioAnalyzer. Libraries were sequenced on Hiseq 4000 with 100 bp paired-end reads.

### RNA Preparation and Sequencing

Dissection of proximal and distal of forelimb buds were performed at E11.5 as described above. The dissected limb buds were stored at −80 in Qiagen RNAlater until genotyping was performed. RNA was extracted from two independent embryos and performed in biological duplicate using RNAeasy Plus mini kit (Qiagen 74134). mRNA enrichment, library preparation and flow-cell preparation for sequencing were performed by the IRCM Molecular Biology Core Facility according to Illumina’s recommendations. Sequencing was done on a HiSeq 2500 instrument with a paired-end 50 cycles protocol.

### ChIP-seq and ATAC-seq Data analysis

ChIP-seq and ATAC–seq reads were aligned to the mm10 genome using bowtie v.2.3.1 with the following settings: bowtie2 -p 8 --fr --no-mixed --no-unal –x. Sam files were converted into tag directories using HOMER v4.9.1 (ref^28^) and into bam files using Samtools v1.4.1 (ref^29^) view function. Peaks were identified by comparing each sample to its input using MACS v2.1.1.20160309 (ref^30^) callpeak function using the parameters: --bw 250 -g mm --mfold 10 30 -p 1e-5. For HOXA11 ChIP, peaks found in both replicates with a pvalue <10^−20^ were considered for further analysis. For Hox13 peaks, HOXA/D13 peaks underwent the same stringent filter. To obtain a high confidence list of specific peaks we considered binding as a specific peak when it did not overlap any peak with a pvalue of 10^−5^ from either replicate of the compared ChIP. This strategy likely eliminate weakly specific or enriched binding (false negative) but allowed to focus on highly pure lists (true positive) of common versus specific peaks.

Heatmaps and average profiles were generated using the Easeq software^31^. ChIP-seq and ATAC-seq data were visualized on the IGV software^32^ using BigWig files generated using the makeUCSCfile HOMER command. For ATAC-seq differential analysis, peaks from ATAC datasets (proximal wild type and distal wild type for the proximo-distal comparison; distal wild type and distal *Hox13*-/- for the WT vs *Hox13*-/- comparison) were merged using HOMER v4.9.1 mergePeaks tool to obtain a file with all the unique position from all the ATAC-seq datasets. ATAC-seq signals were quantified in these different datasets using the analyzeRepeats.pl HOMER command and differential accessibility analyses were performed using getDiffExpression.pl with default parameters, which uses Deseq2 (ref^33^) to perform differential enrichment analysis. Peaks showing differential accessibility of more than two folds and an adjusted pvalue smaller than 0.05 were considered differentially accessible.

### Motif Analysis

Motif analysis on ChIP data was performed using a fixed 200bp window around the peak center. Motif analysis on ATAC-seq data was performed on the whole peak window as the aim is to identify any factors associated with chromatin accessibility and these may not be at the center of the ATAC-seq peak. In all cases, HOMER findMotifsGenome command was used to perform de novo analysis against background sequences generated by HOMER that matches GC content. When the motif rank is not shown, only the top scoring motif based on its p-value is shown.

### RNA-seq Data Analysis

Strand specific paired-end reads were aligned to the mm10 reference genome using TopHat2 (ref^34^) with the parameters --rg-library “L” --rg-platform “ILLUMINA” --rg-platform-unit “X” --rg-id “run#many” --no-novel-juncs --library-type fr-firststrand -p 12. The resulted Bam files were converted to tagDirectory using HOMER and BigWig were produced using the makeUCSCfile HOMER command.

RNA reads quantification was performed using HOMER analyzeRepeats.pl commands with the parameters: rna mm10 -count exons -condenseGenes -noadj -strand -.

Differential expression analysis was performed using HOMER getDiffExpression.pl command with the default parameters which performs differential expression with Deseq2.

### Single Cell ATAC-seq Analysis

Sequencing reads were aligned using Cellranger-atac version1.1.0 (from 10xGenomics®) and the resulted sorted bam files were structured into hdf5 snapfiles (Single-Nucleus Accessibility Profiles) using snaptools version 1.4.1. Quality control analysis was performed using SnapATAC 1.0.0. (ref^35^). Cells with logUMI in [3.5-5] and fragment overlapping peaks above 25% were selected. They were cleaned by eliminating bins overlapping with ENCODE Blacklist regions^36^, mitochondrial DNA as well as the top 5% of invariant features (house-keeping gene promoters). Dimensionality reduction was performed using a non-linear diffusion map algorithm^37^ available in the SnapATAC 1.0.0 package to produce 50 eigen-vectors of which the first 26 were selected (see Extended Data Figure) in order to generate K Nearst Neighbor graph with K=15. The clustering was performed using Leiden Algorithm^38^ available in the package leidenalg version 0.7.0 with a resolution of 0.7 and 2 UMAP (uniform Manifold Approximation and Projection version 0.2.3.1) embedding were generated to visualize the data^39^. Fragments originating from the cells belonging to them same clusters were pooled using snaptools 1.4.1 and peak calling was performed using MACS2 version 2.1.2 for each of the clusters^40^.

### Protein Extraction and Western-blot Analysis

Nuclear extracts were performed using pooled forelimb and hindlimb buds at E11.5 from wild type and *Hoxa11*-/- embryos. Western blot was performed using the anti-HOXA11 antibody (1:500) (SAB1304728, Sigma) and the anti-H3 antibody (1:3000) (abcam) was used as loading control.

### Accession Numbers

The ChIP-seq data for HOX13 in distal limb and RNA-seq for wild type and *Hox13*^−/−^ distal limb buds^13^ at E11.5 are available from the NCBI Gene Expression Omnibus repository under the accession numbers GSE81356. Chicken ChIP-seq data of HOXA13 and HOXA11^10^ were obtained from the accession numbers GSE86088.

### Study Approval

All studies with mice described in this article were approved by the Animal Care Committee of the Institut de Recherches Cliniques de Montréal (protocols # 2015-14 and 2017-10).

