## Extended data for "HOX13-dependent chromatin accessibility modulates the target repertoires of the HOX factors"

**Extended Data Figure 1: Validation of HOXA11 antibody specificity**

Western blot of nuclear extracts from E11.5 wild type and *Hoxa11-/-* limb buds. Two replicates are shown. Antibody against HOXA11 (top panel) and Histone H3 (bottom panel) were used.

Extended Data Fig. 1: Validation of HOXA11 antibody specificity

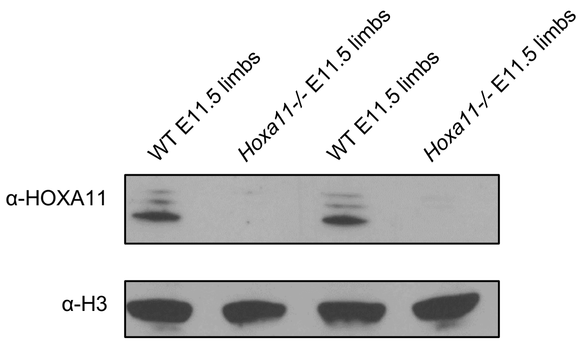

**Extended Data Figure 2: Comparison between HOXA11 and HOXA/D13 binding**

**(a-f)** Venn diagrams showing the number of peak overlaps between the indicated ChIP-seq datasets. Strong peaks refer to peaks with a pvalue $<10^{-20}$. All peaks refers to peaks with an associated pvalue of $<10^{-5}$. Two peaks were considered overlapping whenever the peak window of the two peaks showed any overlap. **(g)** Parameters used to define HOXA11 specific, Common peaks and HOX13-specific peaks in Fig. 1d. **(h)** Genome browser view (IGV) of a few example loci with HOX13-specific (top) and HOXA11-specific (bottom) peaks.

Extended Data Fig. 2: Comparison between HOXA11 and HOXA/D13 binding

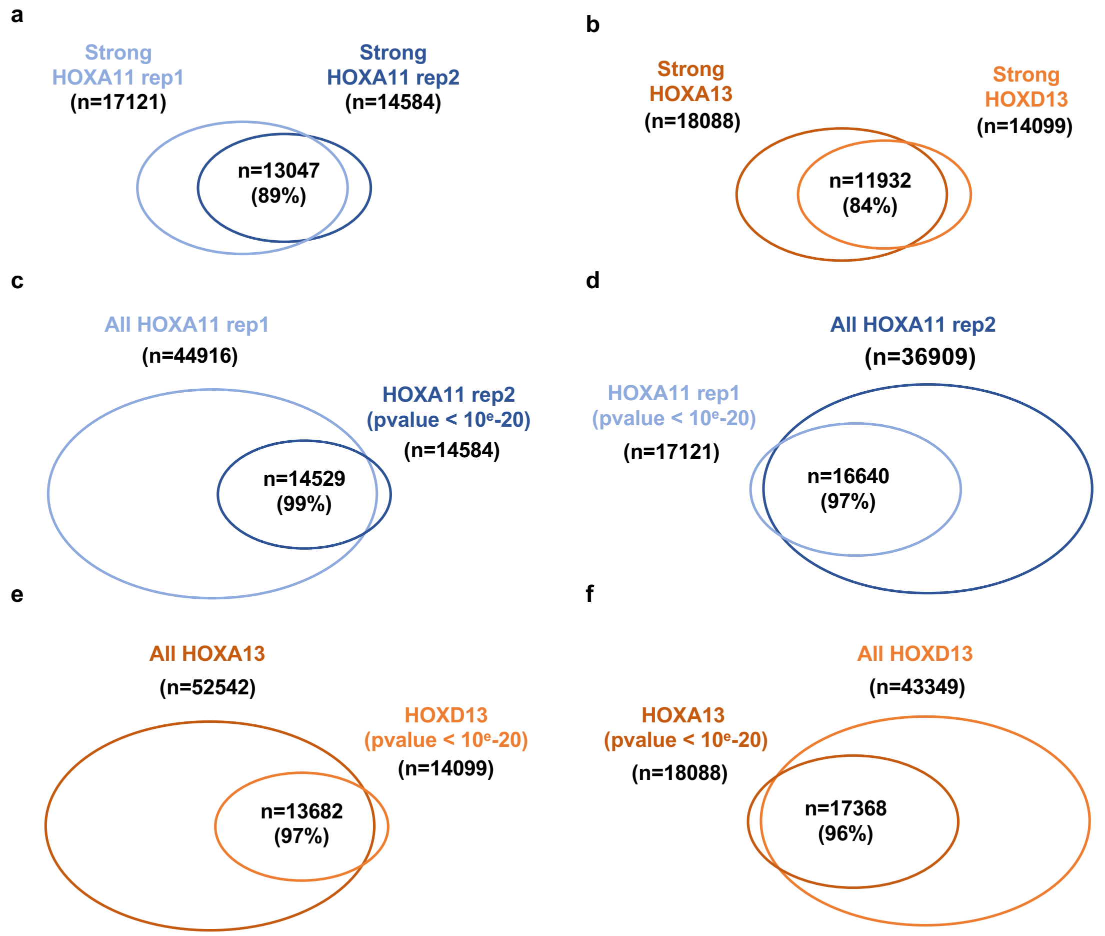


g

| | HOXA13 | | HOXD13 | | HOXA11 rep1 | | HOXA11 rep2 | |
|---|---|---|---|---|---|---|---|---|
| | All | Strong | All | Strong | All | Strong | All | Strong |
| HOXA11 specific: n=2128 | - | - | - | - | + | + | + | + |
| Common HOXA11-HOX13 n=6967 | + | + | + | + | + | + | + | + |
| HOX13 specific: n=2130 | + | + | + | + | - | - | - | - |

h

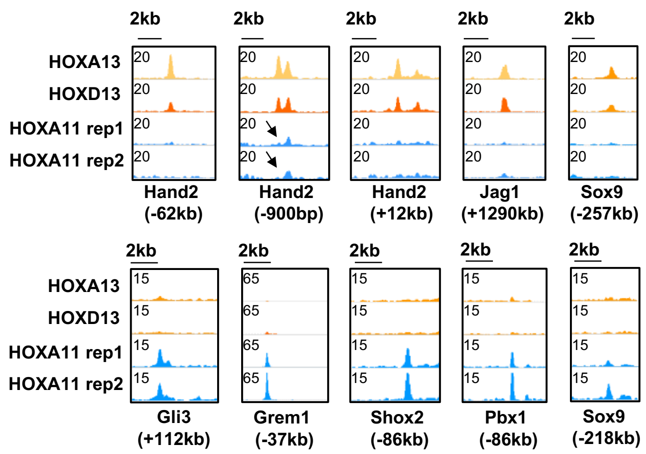

**Extended Data Figure 3: HOXA11 ectopic binding after distal HOXA11 expression**

**(a)** Genome browser view (IGV) of a few example loci with ectopic binding of HOXA11 only when co-expressed distally with HOX13. **(b)** Heatmaps showing a 4kb window of the ChIP-seq signal from the indicated datasets centered on HOXA11 peaks in $R26^{A11d/A11d}$ separated into two clusters using kmeans based on HOX13 signal intensity. **(c)** *De novo* motif analysis (using HOMER) on 200bp windows surrounding the two subsets of ectopic HOXA11 binding. The top scoring motif based on its pvalue is shown.

Extended Data Fig. 3: HOXA11 ectopic binding after distal HOXA11 expression

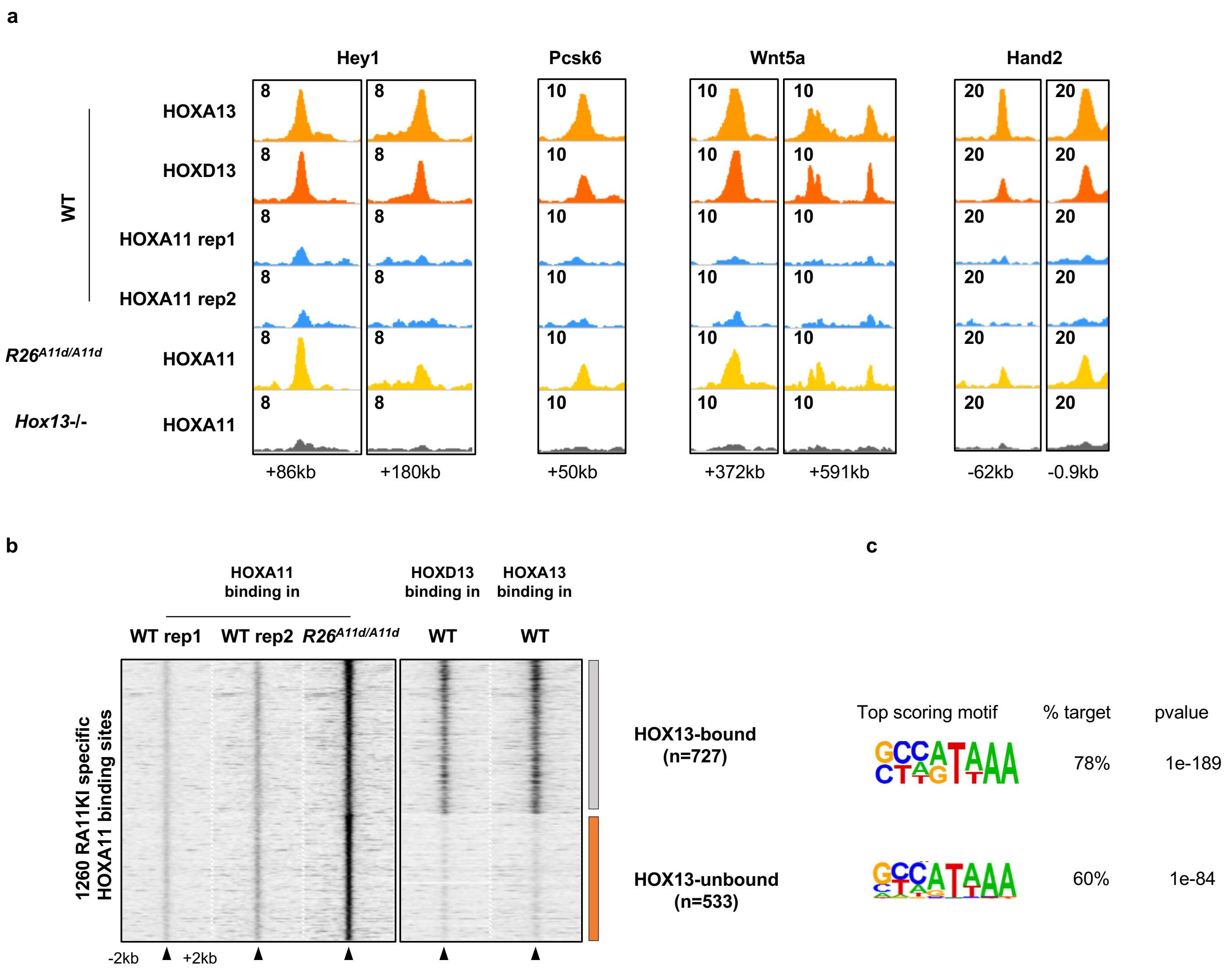

**Extended Data Figure 4: Characterization of proximo-distal E11.5 limbs open chromatin landscapes**

**(a)** Average profile showing proximal and distal ATAC-seq signals at proximal enriched (left) and distal enriched (right) ATAC-seq peaks. **(b)** *de novo* motif analyses (HOMER) at proximal enriched ATAC peaks using the whole open region as input rather than a fixed window. **(c)** GREAT analysis showing the enriched GO terms for biological processes for genes associated with proximal (top, blue) and distal (bottom orange) ATAC-seq peaks. **(d)** Volcano Plot showing the log2 fold change of proximal over distal RNA expression (on the X axis) and the adjusted pvalue computed by DESEQ2 differential expression analysis (on the Y axis). Proximally enriched genes are shown in blue while Distally enriched genes are in orange. **(e)** Gene ontology associated with distally enriched genes. **(f)** Boxplot showing the distance between the TSS of distal enriched genes and the closest proximal- (left) or distal- (right) enriched ATAC-seq peak. Center lines show medians; box limits indicate the twenty fifth and seventy-fifth percentiles; whiskers extend to 1.5 times the interquartile range from the twenty-fifth to seventy-fifth percentiles. p-value was calculated using unpaired two-tailed student t-test.

### Extended Data Fig. 4: Characterization of proximo-distal E11.5 limbs open chromatin landscapes

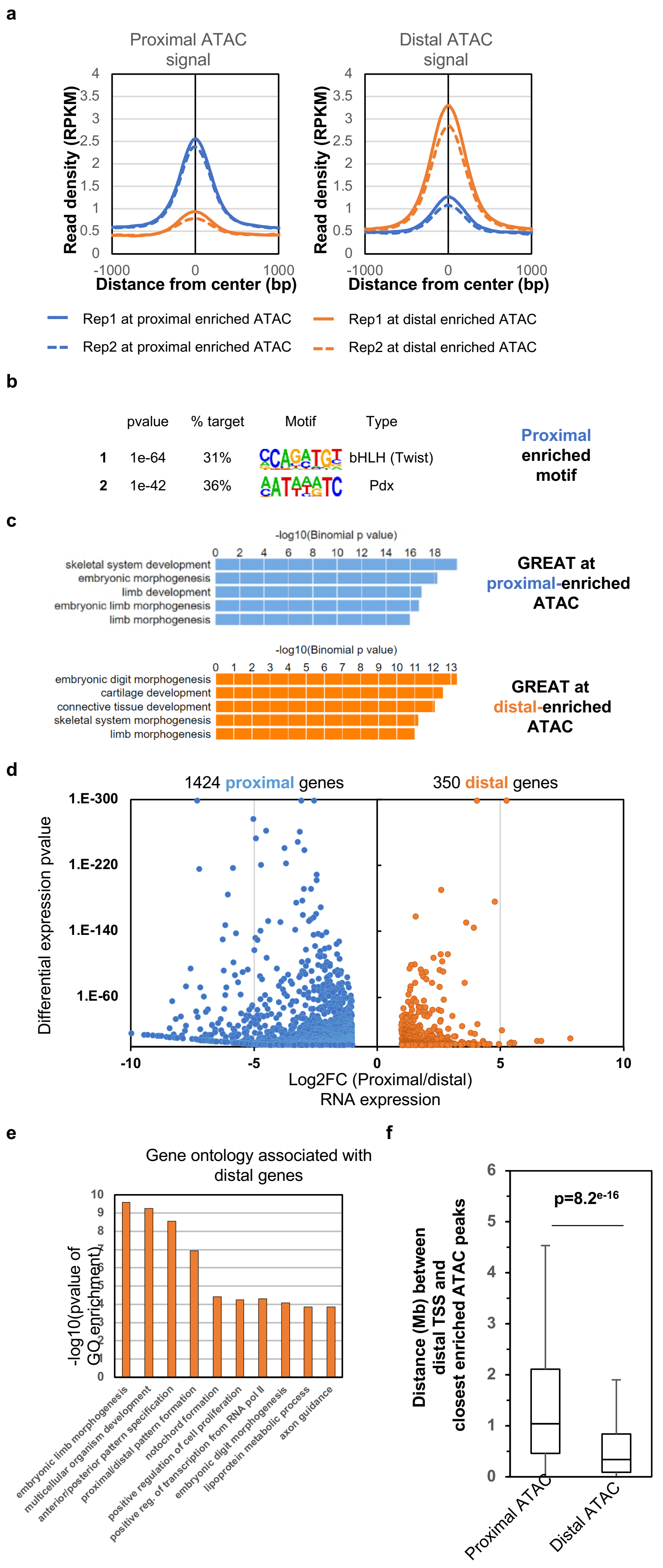

**Extended Data Figure 5: Analysis of HOX13-dependent open chromatin landscape**
**(a)** Dispersion plot showing the correlation between the replicates of wild type (left) and *Hox13*-/- (right) ATAC-seq signal (reads per million reads per kb). Pearson correlation is shown. **(b)** Boxplot showing HOXA13 (left) and HOXD13 (right) binding intensity (reads per million read per kb) in a +/-200bp window surrounding the indicated ATAC peak category. Center lines show medians; box limits indicate the twenty fifth and seventy-fifth percentiles; whiskers extend to 1.5 times the interquartile range from the twenty-fifth to seventy-fifth percentiles. **(c)** Heatmaps showing a 4kb window of the indicated ATAC-seq signal intensity distal specific ATAC-seq peaks clustered using k-means into two clusters based on the ATAC signal in *Hox13*-/-. **(d)** *De novo* motif analysis (HOMER) showing the top scoring motif (based on its pvalue) enriched in a 200bp window at the distal sites showing Hox13-independent and Hox13-dependent accessibility. **(e)** Dispersion plot showing the HOXA13 (X axis) and HOXA11 (Y axis) signals at the indicated ATAC-seq peaks categories. **(f)** Boxplot showing the log2 fold change of WT over *Hox13*-/- differential RNA expression for proximal-enriched, unchanged and distal-enriched genes. Center lines show medians; box limits indicate the twenty fifth and seventy-fifth percentiles; whiskers extend to 1.5 times the interquartile range from the twenty-fifth to seventy-fifth percentiles. pvalue were calculated using unpaired two-tailed student t-test. **(g)** Genome browser view showing the indicated ChIP-seq and ATAC-seq datasets at a few example loci that undergo loss of accessibility in *Hox13*-/- distal E11.5 limb buds. **(h)** Heatmap showing wild type and *Hox13*-/- RNA expression values at distal enriched genes (from GSE81358). Rows are centered; unit variance scaling is applied to rows. Rows are clustered using Euclidean distance and average linkage. 361 rows, 4 columns.

Extended Data Fig. 5: Analysis of HOX13-dependent open chromatin landscape

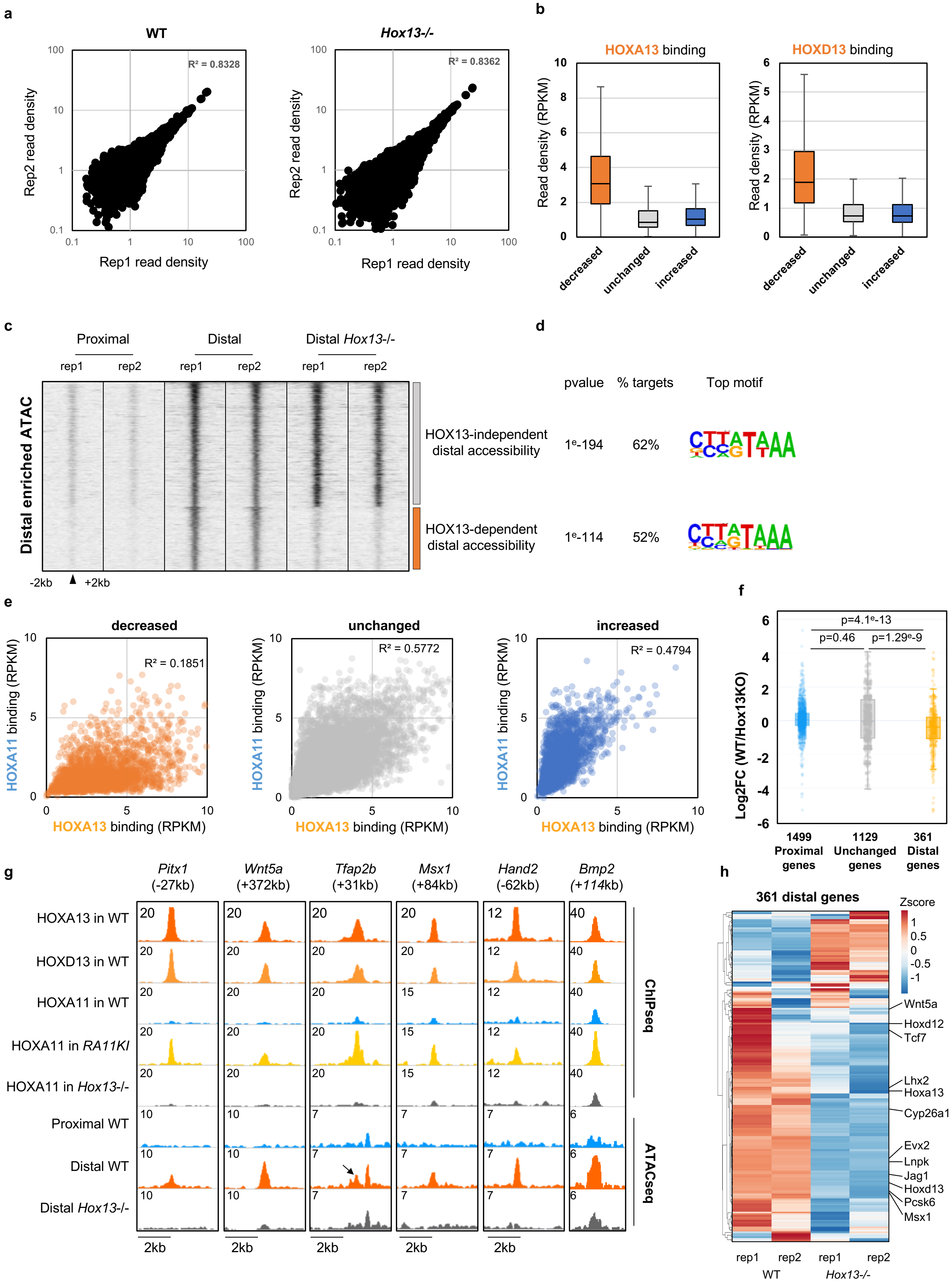

**Extended Data Figure 6: Pipeline and quality control of scATACseq on WT and *Hox13*-/- limb bud**

(**a**) Scheme of the single-cell ATAC-seq assay and data analysis performed. **(b)** Description of the pipeline used to analyze the single-cell ATAC-seq datasets generated in this study. **(c-d)** Dispersion plot of wild type **(c)** and *Hox13*-/- **(d)** showing the log10 (UMI number per each barcode) on the X axis and the fraction of reads in peaks per barcode on the Y axis. The red square shows the threshold that we used to consider barcodes as representing a captured cell. **(e)** Pairwise comparison of dimensions to determine the number of dimensions to use for dimensionality reduction (UMAP) and clustering of the scATAC-seq datasets.

Extended Data Fig. 6: Pipeline and quality control of scATACseq on WT and *Hox13*-/- limb bud

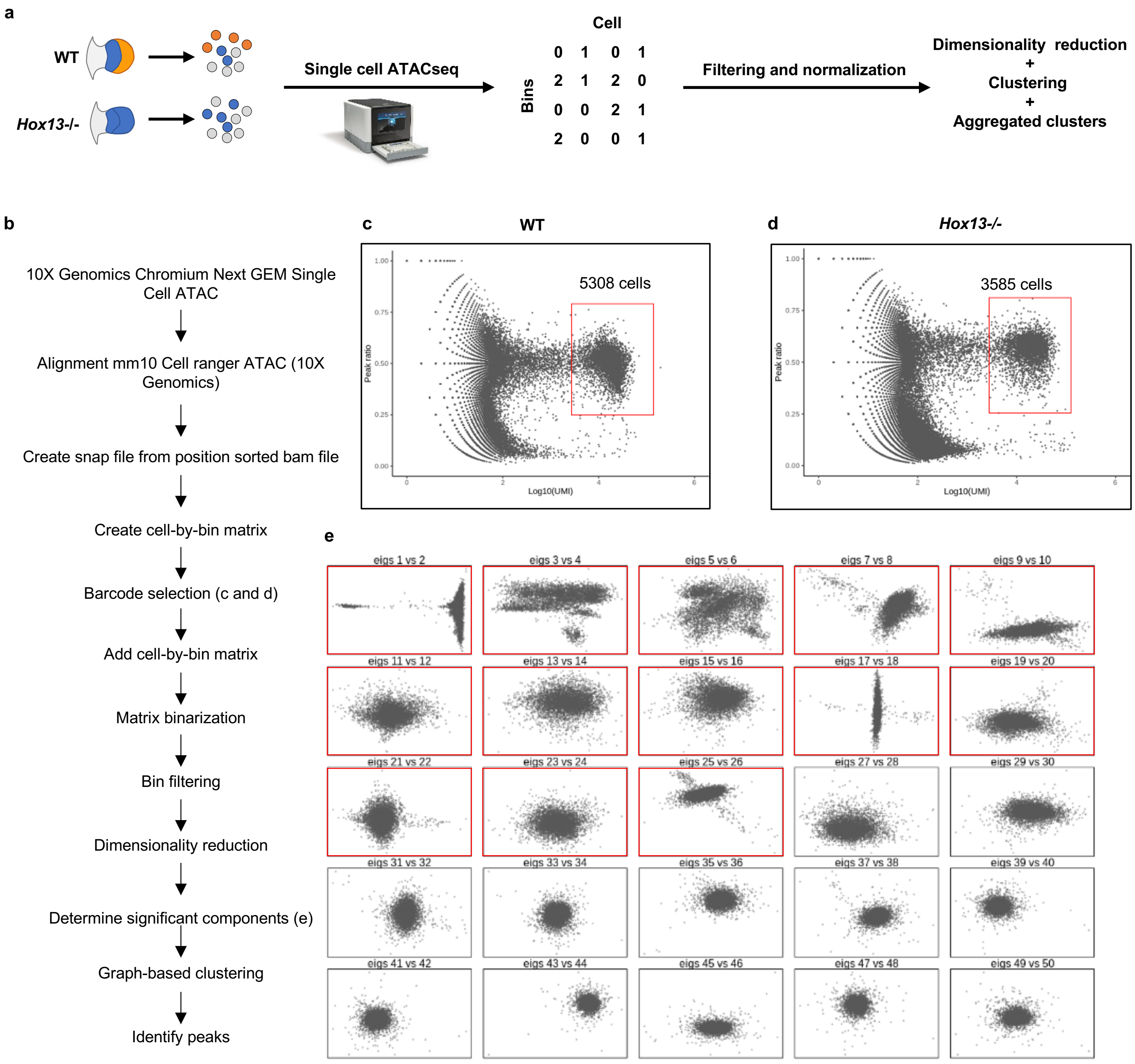

**Extended Data Figure 7: Annotation of clusters identified by single cell-ATACseq**

Genome browser view (IGV) of aggregated signal of single cell ATAC-seq clusters separated by genotype (wild type, colored and *Hox13*-/-, in grey) at marker loci for the indicated markers. Other markers were used to confirm correct identification (not shown).

Extended Data Fig. 7: Annotation of clusters identified by single cell-ATACseq

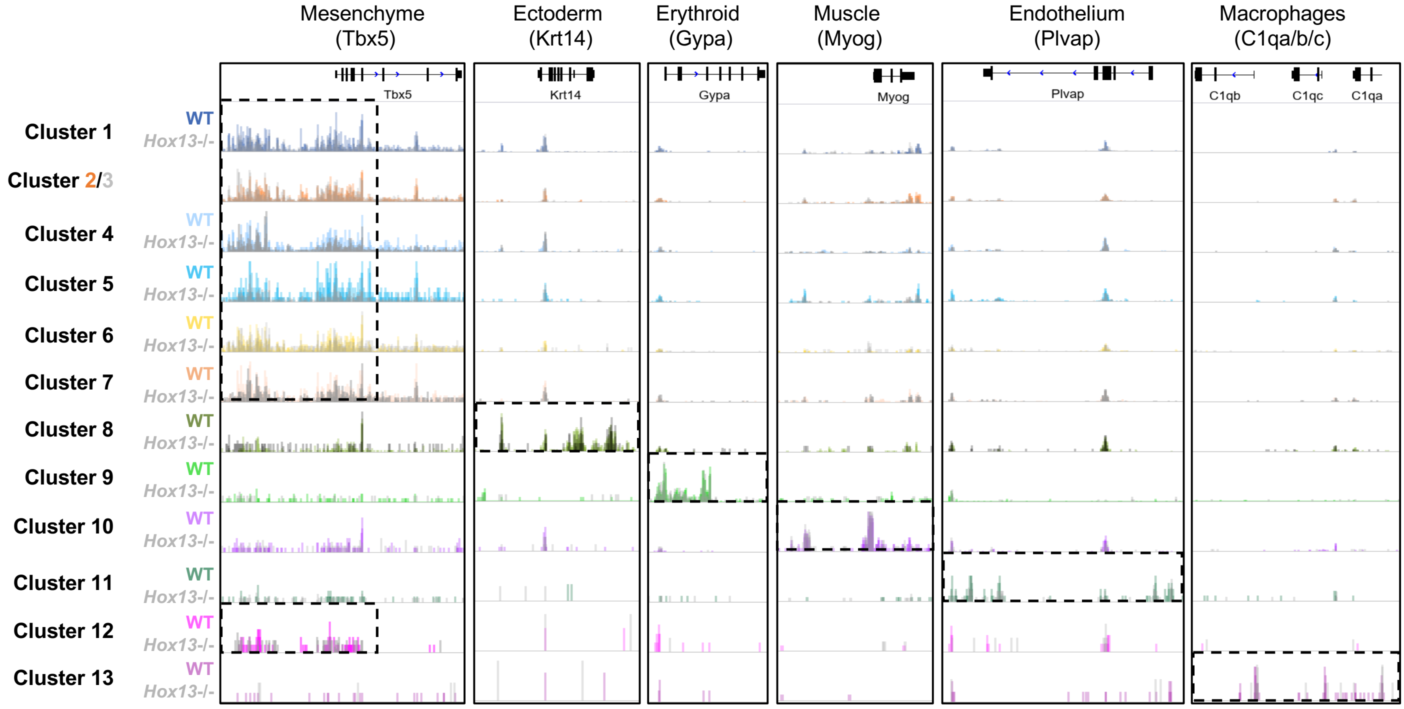

**Extended Data Figure 8: Non-mesenchymal lineages are weakly affected by Hox13 loss of function**

**(a)** Dimensionality reduction map (UMAP) colored by genotype (wild type in orange, and *Hox13*-/- in grey). **(b)** Magnification of the squares from **(a)** separated by genotype (wild type, top and *Hox13*-/-, bottom) showing the UMAP coordinates from the non-mesenchymal lineages. **(c)** Bar plot showing the percentage of each mesenchymal (left) and non-mesenchymal clusters from wild type (orange) and *Hox13*-/- (grey).

Extended Data Fig. 8: Non-mesenchymal lineages are weakly affected by Hox13 loss of function

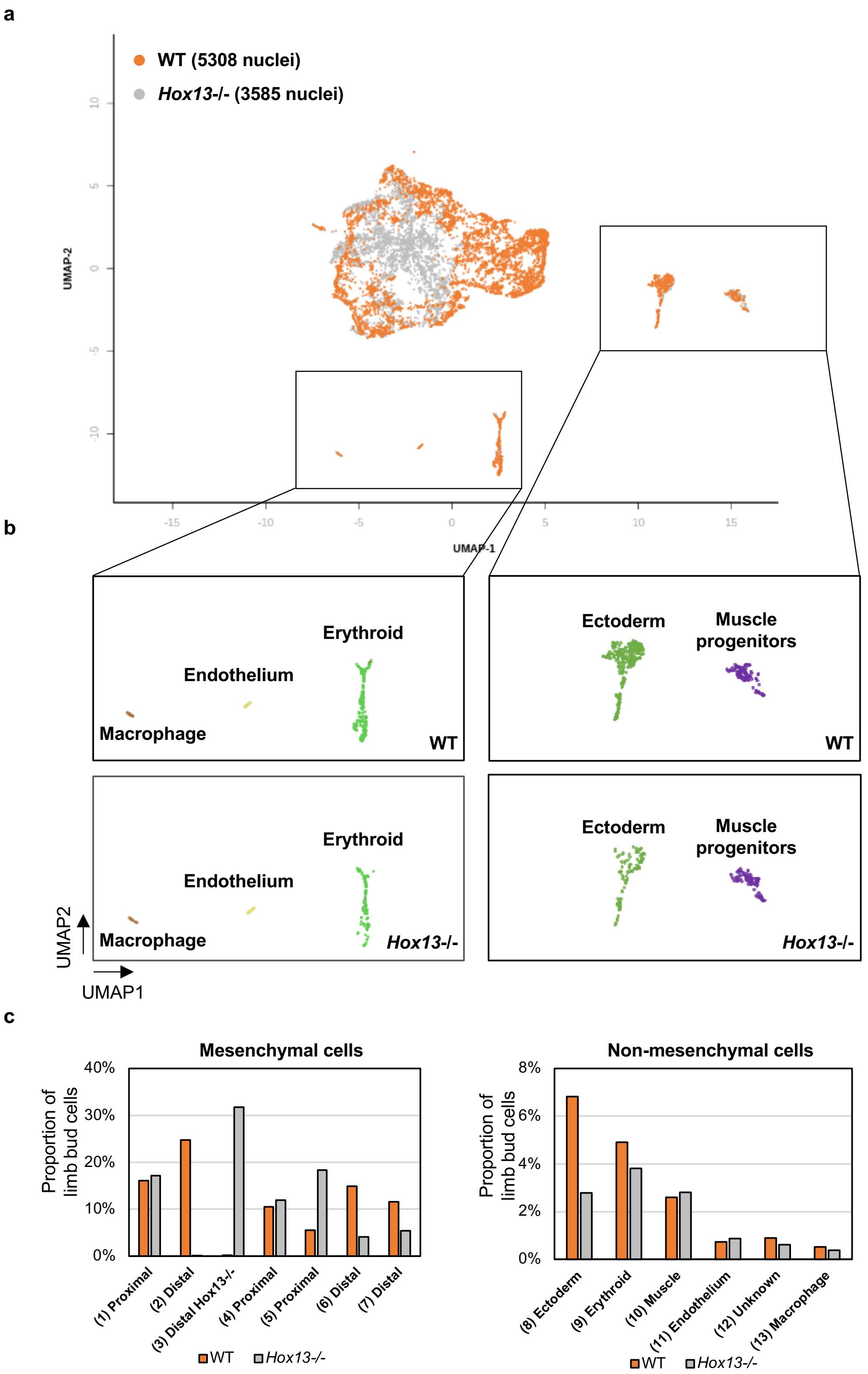

**Extended Data Figure 9: Opening of the enhancer driving *Hoxa11* antisense transcription is conserved in Chicken**

**(a)** Pairwise sequence conservation at the *Hoxa*11 locus. The HOX13-dependent enhancer driving antisense transcription in distal mouse limb bud is highlighted. **(b)** Genome browser view (IGV) of HOXA13 and HOXA11 ChIP-seq data (GSE86089) from primary chicken mesenchymal limb progenitor cells and ATAC-seq from chicken wing bud at early stage (HH20, blue), prior *Hox13* expression and late (HH26/27, orange) distal wing bud, in the *Hox13*-expressing domain at the *Hoxa11* locus.

Extended Data Fig. 9: Opening of the enhancer driving *Hoxa11* antisense transcription is conserved in Chicken

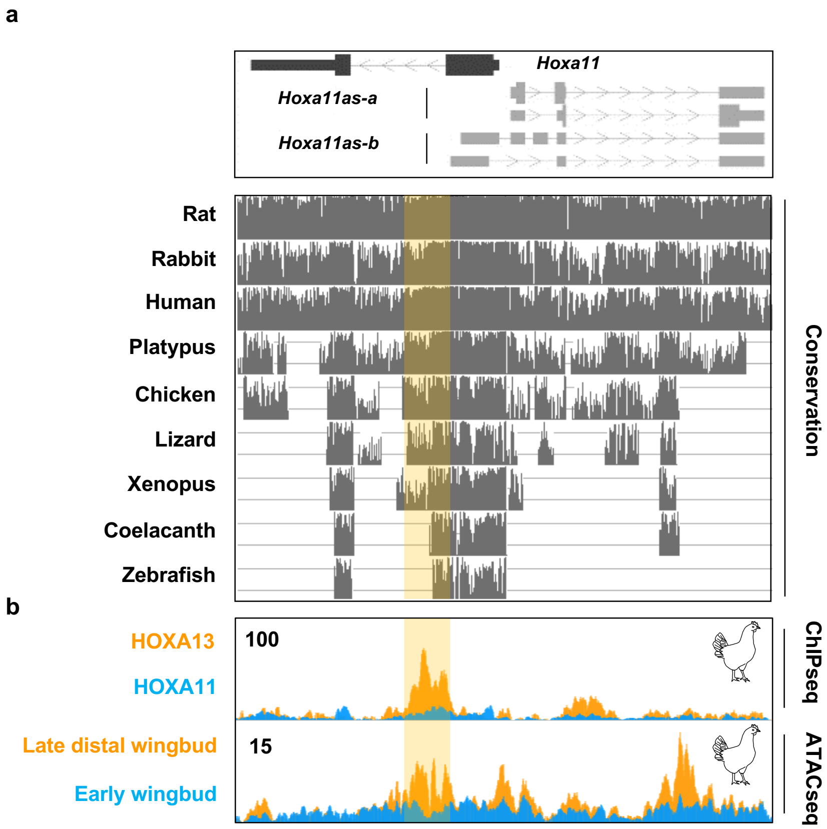
